# Self-Assembling Peptide Hydrogels - PeptiGels^®^ as a Platform for Hepatic Organoid Culture

**DOI:** 10.1101/2021.03.01.433333

**Authors:** Adedamola Olayanju, Aline F Miller, Tahera Ansari, Christopher E. Goldring

**Affiliations:** Manchester BIOGEL, Mereside, Alderley Park, Alderley Edge, Cheshire, SK10 4TG, UK; MRC Centre for Drug Safety Science, Department of Molecular and Clinical Pharmacology, Institute of Translational Medicine, University of Liverpool, Liverpool, United Kingdom, L69 3GE; Faculty of Medical Science, Division of Surgery and Interventional Science, UCL, Gower St., London WC1E 6BT; Tissue Engineering and Regenerative Medicine, Northwick Park Institute for Medical Research (NPIMR), Harrow, United Kingdom, HA1 3UJ

**Keywords:** 3D cultures, Organoids, Extracellular Matrix, Peptide Hydrogels, PeptiGels^®^

## Abstract

A major challenge in advancing preclinical studies is the lack of robust in vitro culture systems that fully recapitulate the *in vivo* scenario together with limited clinical translational to humans. Organoids, as 3-dimensional (3D) self-replicating structures are increasingly being shown as powerful models for *ex vivo* experimentation in the field of regenerative medicine and drug discovery. Organoid formation requires the use of extracellular matrix (ECM) components to provide a 3D platform. However, the most commonly used ECM, essential for maintaining organoid growth is Matrigel and is derived from a tumorigenic source which limits its translational ability. PeptiGels^®^ which are self-assembling peptide hydrogels present as alternatives to traditional ECM for use in 3D culture systems. Synthetic PeptiGels^®^ are non-toxic, biocompatible, biodegradable and can be tuneable to simulate different tissue microenvironments. In this study, we validated the use of different types of PeptiGels^®^ for porcine hepatic organoid growth. Hepatic organoids were assessed morphologically and using molecular techniques to determine the optimum PeptiGel^®^ formulation. The outcome clearly demonstrated the ability of PeptiGel^®^ to support organoid growth and offer themselves as a technological platform for 3D cultured physiologically and clinically relevant data.

## Introduction

A major challenge in translating preclinical studies is the lack of robust *in vitro* culture systems that fully recapitulate the *in vivo* scenario. Most current models lack direct translational relevance to humans e.g. rodents hinder clinical adoption of emerging therapies due to the possibility of severe adverse reactions. This is particularly evident with drug induced liver injury (DILI) which accounts for more than 50% of acute liver failure (1). Although studies conducted in small rodent hold value in the understanding of fundamental concepts, the inability to fully extrapolate results to clinical findings has had a major impact in the acceleration of regenerative medicine, cell therapy and drug discovery.

Organoids, as 3-dimensional (3D) self-replicating structures are increasingly being shown as powerful models for *ex vivo* experimentation due to their ability to recapitulate and maintain physiological phenotypes of their tissues of origin (2). Organoids can overcome several limitations seen with 2D cultures such as de-differentiation and lack of cell-ECM communication leading to altered phenotypic expression. Additionally, the use of organoids may help to reduce the number of animals used in research, whilst still providing physiologically and clinically relevant data

Organoid formation requires the use of extracellular matrix (ECM) as a platform for the 3D conformation. Most ECMs used are animal-derived and have shown several limitations such as batch-to-batch variability, lack of tuneability and ease of use. In particular, Matrigel the most common, is derived from a tumorigenic source limiting its translational ability (3). Hence, there is a critical need for a synthetic alternative ECM in the generation of organoids.

Over recent years, significant resources have been directed at designing 3D scaffolds that mimic the ECM platforms including self-assembling peptides that can be hydrated to mimic the cellular niche (4). Of particular interest are synthetic and biologically relevant PeptiGel^®^, which are non-toxic, biocompatible, and biodegradable. Their tuneable properties make them attractive platforms to simulate different tissue microenvironments.

An increasing number of studies have demonstrated the potential use of synthetic self-assembling peptides in culturing 3D cells to create physiologically and reproducibly relevant *in vitro* models that recapitulates the *in vivo* counterpart (4, 5). Self-assembling peptide hydrogels such as PeptiGels^®^ are hydrogels made from short amino acid amphipathic peptides that self-assemble into β-sheet forming structures (6). Above a critical concentration, there is a formation of transparent hydrogel. Importantly, these peptide hydrogels are tuneable to give hydrogels of different mechanical and functional properties, hence having the ability to replicate different tissues of interest (6). Increasing evidence have shown that these self-assembling peptides are biocompatible and support the growth and propagation of different cell types in 3-dimensional conformation and have supporting roles to drive cell differentiation, model diseases and show potentials for other investigative biological functions (5, 7). The adoption of alternative synthetic ECM sources such as synthetic self-assembling peptide hydrogels PeptiGels^®^ in the field of 3D cell culture may produce physiologically and clinically relevant data. Therefore, the aim of this study was to validate the use of PeptiGels^®^ for the growth of organoids and to explore which formulation of PeptiGel^®^ offers the optimal environment for hepatic organoids.

## Materials and Methods

The use of Manchester BIOGEL PeptiGels^®^ as an alternative source of ECM for the growth and maintenance of porcine hepatic organoids was investigated. All experiments were conducted in accordance with the Home Office Regulation on animal tissue usage.

Approx. 1 g of liver tissue was harvested from pigs undergoing termination from unrelated studies at NPIMR. A mixture of single cells were isolated from healthy porcine liver tissue and propagated (2) on five PeptiGels^®^ (Alpha 1-5); each with a different mechanical property and/or charge (see table 1) with the aim of determining the optimum cell culture parameters. The resulting organoids were assessed for their 3D morphology using light microscopy techniques and compared to organoids grown on conventional ECMs i.e. Matrigel as it is the current gold standard in organoids generation and therefore used as a control. Matrigel and PeptiGel^®^ generated organoids were lysed in RLT buffer and using Qiagen’s RNeasy Mini Kit as instructed by the manufacturer was employed to extract RNA from Matrigel and PeptiGel^®^-generated organoids. The cDNA was synthetized using Promega’s GoScript Reverse transcription systems as instructed by the manufacturer. The resulting Organoids cDNA were phenotyped by Real-time quantitative PCR for Cytokeratin-18 (CK18) and Gapdh gene expressions using Biorad CFX100 PCR instrument.

**Table 1.**
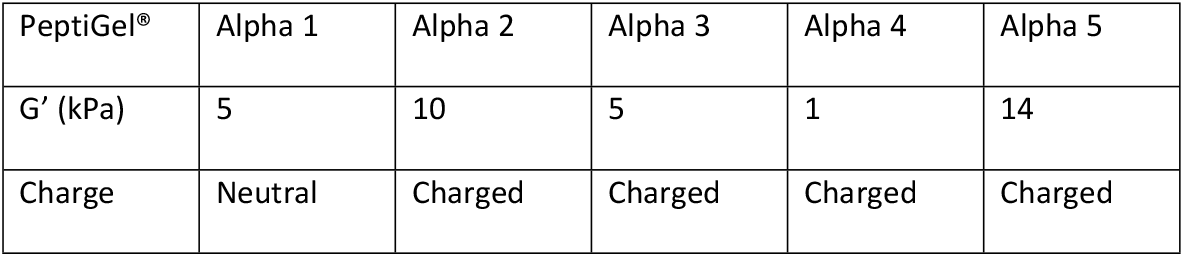
The mechanical properties of different versions of PeptiGels^®^

| PeptiGel® | Alpha 1 | Alpha 2 | Alpha 3 | Alpha 4 | Alpha 5 |
| --- | --- | --- | --- | --- | --- |
| G' (kPa) | 5 | 10 | 5 | 1 | 14 |
| Charge | Neutral | Charged | Charged | Charged | Charged |

## Results

### Time course generation of hepatic organoids using PeptiGel^®^ Technology

PeptiGel^®^-generated organoid units were maintained in liver organoid media as described by Broutier *et al* (2). The seeded cells initially remained as single cells (days 1-2), but over time, cells started to coalesce to form the 3D structures (organoids). By day 6, they showed the classic organoid-like appearance involving the formation of a 3-dimensional structures with clusters of cells (Fig 1). Notably, hepatic cells grown in PeptiGel^®^ Alpha 5 showed visible organoids by day 6. By day 13, they were fully formed organoids within the cultures with PeptiGels^®^ 1, 2 and 3. An assessment of the organoids demonstrated that the hepatic cells showed different morphologies when grown on different versions of the PeptiGels^®^ when compared to the control organoids grown on Matrigel (Fig. 1). PeptiGel^®^-generated organoids using Alpha 1 showed organoid like structures by day 14 but by day 27, they had fully dissociated. PeptiGel^®^-generated organoids using PeptiGels^®^ Alpha 2-4 showed similar morphologies to the those grown on PeptiGel^®^ Alpha 1 but those generated using PeptiGel^®^ Alpha 2 showed a higher rate of formation compared to those grown on PeptiGels^®^ Alpha 3 and Alpha 4. (Fig. 1). Notably, organoids generated using PeptiGel^®^ Alpha 2 were similar to those grown using Matrigel.

**Figure 1:**
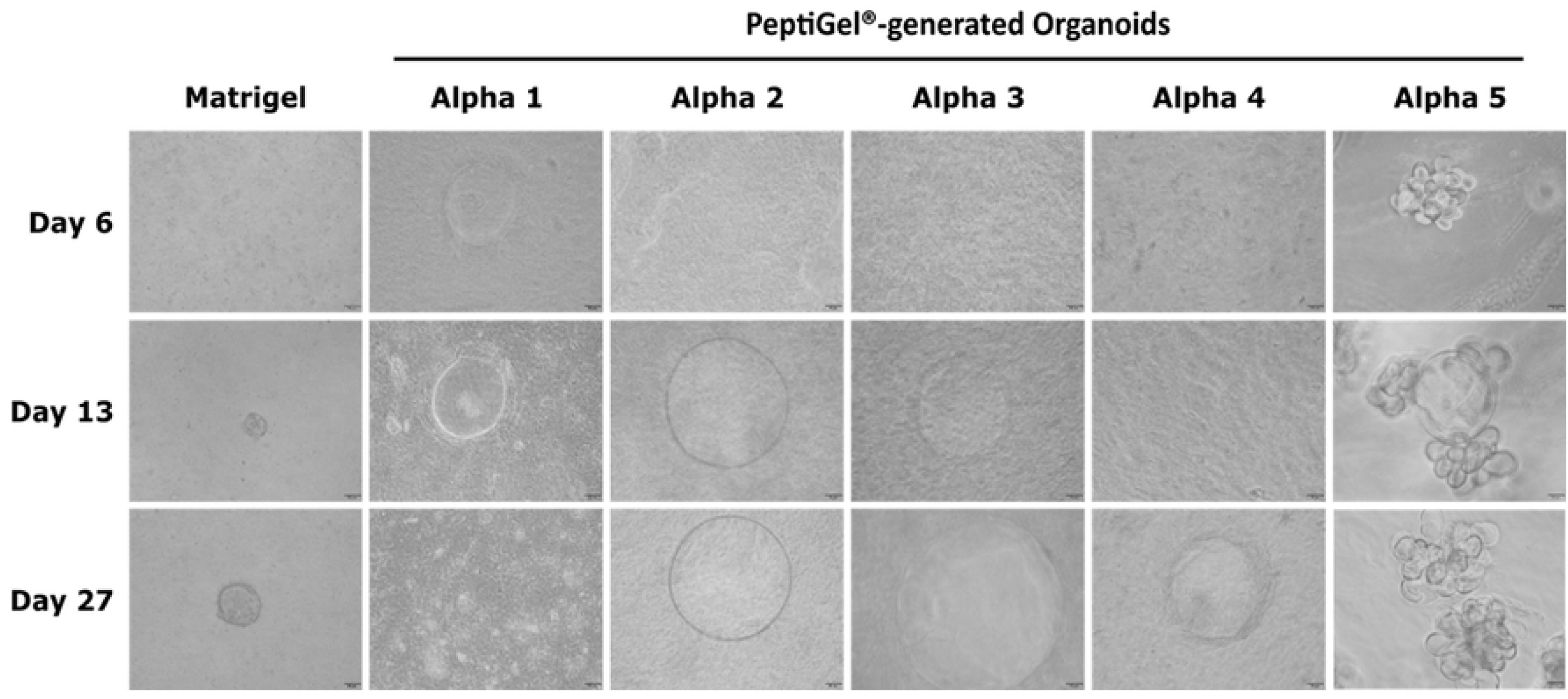
PeptiGel^®^-generated hepatic organoids. Time course establishment of porcine liver organoids using PeptiGels^®^ technology platform. Scale bar is 100 μm.

### RNA extraction and PCR analysis of Cytokeratin-18 genes from PeptiGel^®^-generated organoids

Having established that self-assembling peptide hydrogels such as PeptiGels^®^ offered a platform for generating 3D organoids, we assessed whether PeptiGel^®^-generated organoids were compatible with current methods for RNA isolation, specifically to assess if there was any potential interaction between RNA in the organoids and the biomaterial and to validate that the organoid were in fact liver specific organoids.

RNA was extracted and detected in the organoids generated from PeptiGels^®^ but was lower when compared to RNA extracted from Matrigel (Fig. 2A). When comparing the organoids grown in PeptiGels^®^, PeptiGel^®^ Alpha 2 organoids had the highest RNA concentration followed by PeptiGel^®^ Alpha 4 organoids. The high values in PeptiGel^®^ Alpha 2 organoids could be due to the higher expression of organoids seen with the cultures (Fig. 1). PeptiGels^®^ Alpha 3, 1 and 5 had the lowest values respectively and may be due to the low formation of the organoids in these cultures and/or the formation of peptide fibrils between the gel and the RNA.

**Figure 2A:**
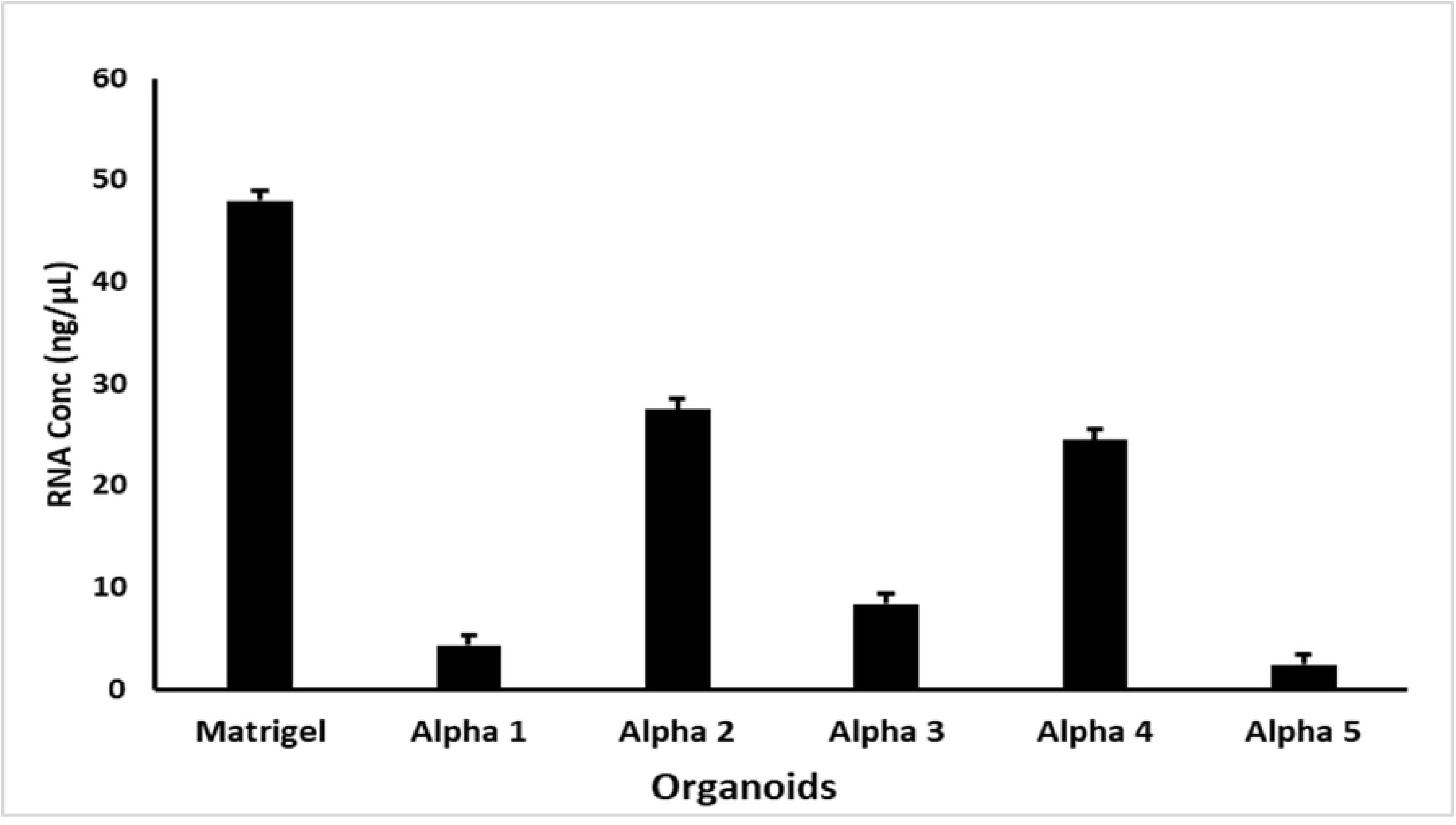
RNA Measurements in PeptiGel^®^ and Matrigel-organoids. Measurements were taken using Nanodrop ND1000, Thermo Scientific, Wilmington, USA. All data represent the mean + SD of n=3 samples.

Real-time quantitative PCR analysis of Cytokeratin-18 (CK18) gene and Gapdh was performed to check if the RNA extracted from the PeptiGel^®^-organoids were compatible with common genetic analysis such as PCR and if they expressed cell markers. For primer sequences, see supplementary table 1. All PeptiGel^®^-organoids express CK18, a marker of epithelial cells (Fig. 2B). They also expressed Gapdh as a loading control. Although, there were low expression of these genes in these samples which could be due to the initial RNA normalisation of all samples to the sample with the lowest RNA value (PeptiGel^®^ Alpha 5-organoid) which means the starting RNA value to be synthesized was low. However, these genes were still shown to be expressed indicating the presence of epithelial cells in all the PeptiGel^®^ cultures.

**Figure 2B:**
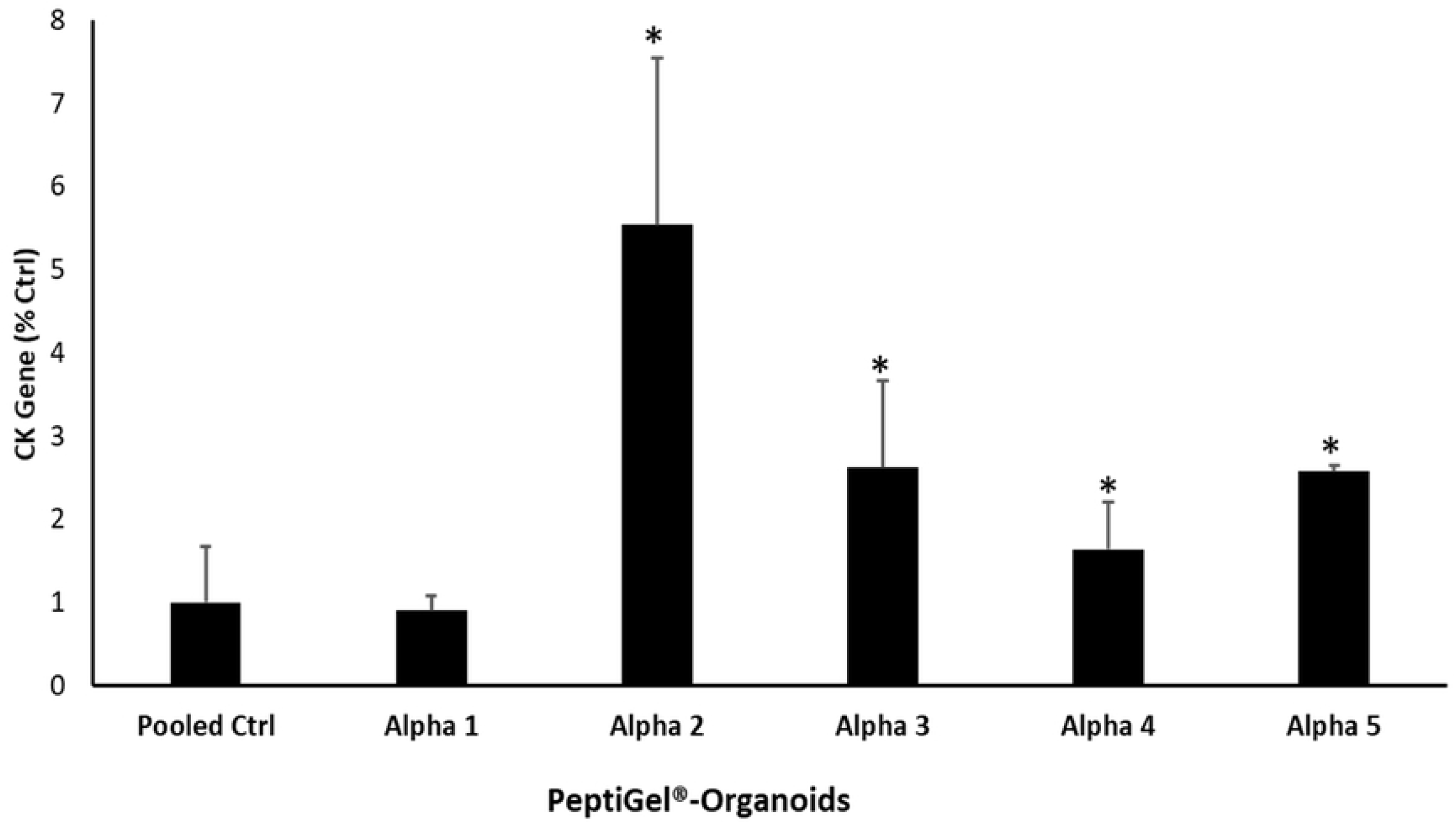
PeptiGel^®^-organoids express epithelial cell marker CK18 gene. Real-time quantitative PCR analysis of CK18 gene in PeptiGel^®^ generated organoids. CK18 gene level was normalised to the loading control, Gapdh, and expressed as a percentage of CK18 level of the pooled control. All data represent the mean + SD of *n*=3 experiments. Statistical analyses were performed using an unpaired *t* test (\**P*<0.05 vs pooled control).

## Discussion

PeptiGels^®^ were assessed to determine their potential ability to support the growth and development of organoids with appropriate morphology and the expression of tissue-specific markers. We have shown that it is possible to use PeptiGels^®^ as platforms to generate organoids derived from porcine hepatic cells. In particular, PeptiGel^®^ Alpha 2 generated organoids with a classic hepatic organoid phenotype and interestingly the mechanical properties of PeptiGel^®^ Alpha 2 is G’ prime of 10kPa and correlates to that seen in diseased liver tissue (8). PeptiGel^®^ Alpha 5 generated organoids showed more branching when compared to the other organoids. This change in morphology may be have been influenced by the mechanical property of the gel (14kPa). As such, this may represent a suitable *in vitro* model for investigating the dynamics associated with fibrotic livers, where the tissue is stiffer than normal. The present investigation needs to be expanded to confirm that the organoids generated using PeptiGel^®^ behave as expected with regards to cryopreservation, the passaging of the organoids, and the growth of organoids for an extended period of time. In addition, more robust internal cellular morphological analysis, differentiation and spatial arrangement of cells, proteomic and genotypic phenotyping and how these parameters evolve with time requires further investigation. Furthermore, the use of functionalised PeptiGels^®^ such as in the incorporation of Integrin peptide motifs (RGD), Laminin peptide motifs (IKVAV, YIGSR) or Collagen peptide motif (GFOGER) with variable mechanical strengths will be explored to fine-tune the generation of hepatic organoids.

The recapitulation of the *in vivo* counterpart by cells grown in 3D matrices requires an analysis of the genetic or protein responses in the matrices; analysis of which requires good quality RNA or protein for further downstream PCR analysis. Hence, a preliminary phenotyping of the PeptiGel^®^-generated organoids using PCR was carried out to assess the presence of tissue-specific markers. RNA was successfully extracted using a column-based approach, however, the yield observed in this study was low compared to the extraction in Matrigel-generated organoids (Fig. 2A). This reduction in the yield was probably due to the presence of fibrils. A study by Burgess *et al.,* 2018 (9) undertook a comparative analysis of RNA extraction from cells encapsulated in self-assembling peptide hydrogels using two different extraction principles: solution-based extraction and direct solid-state binding of RNA and showed the latter was a more efficient method of extraction (9). In line with this study, we have shown that the use of a direct solid-state binding of RNA extraction method allowed the extraction of RNA of PeptiGel^®^-organoids. Burgess *et al.,* also showed that the use of pronase, a broad-spectrum enzyme solution reduced the amount of fibril present and increased RNA yield (9). Therefore, future investigation will aim to increase the yield of the RNA content by employing the use and impact of pronase to digest the organoids lysate before being subjected to RNA analysis.

cDNA was synthesized from the extracted RNA from PeptiGel^®^-generated organoids and the expression of CK18 (epithelial cell marker) and Gapdh (housekeeping) genes was carried out using PCR. CK18 and Gapdh was expressed in all PeptiGels^®^ platforms used (Fig. 2B).The low expression of these genes seen may in part be due to the low yield of RNA content and initial RNA normalisation of all samples to the sample with the lowest RNA value (PeptiGel^®^ Alpha 5-organoid) which means the starting RNA value to be synthesized was low. However, these genes were still shown to be expressed and their expression could be improved upon by increasing the starting sample RNA concentration.

We believe this is the first report to show that self-assembling peptides-PeptiGels^®^ can be used for the generation of hepatic organoids. Organoids have been shown to be powerful preclinical models to accelerate clinical developments. However, organoids generated using animal-derived ECM have limitations in areas of safety and ethical use. PeptiGel^®^-generated organoids may therefore enhance translational research. This is envisaged to have a substantial impact in the development of regenerative medicine and accelerate drug discovery processes. Our present findings show the potential of PeptiGel^®^ technology platform as a suitable alternative to ECM required for organoid culture and as such, may have huge applications in the development of cellular therapeutic diagnostic tool, translation of *in vitro* data to appropriate and robust preclinical studies and eventual clinical application.

## Acknowledgement

The authors are grateful for the financial support received from 3DBioNet Collaborative grant, the donation of PeptiGels^®^ from Manchester BIOGEL and porcine tissue from NPIMR.

## Abbreviations

3D: 3-dimensional
DILI: Drug Induced Liver Injury
ECM: Extracellular Matrix
CK18: Cytokeratin-18

---

**Supplementary Table 1:**
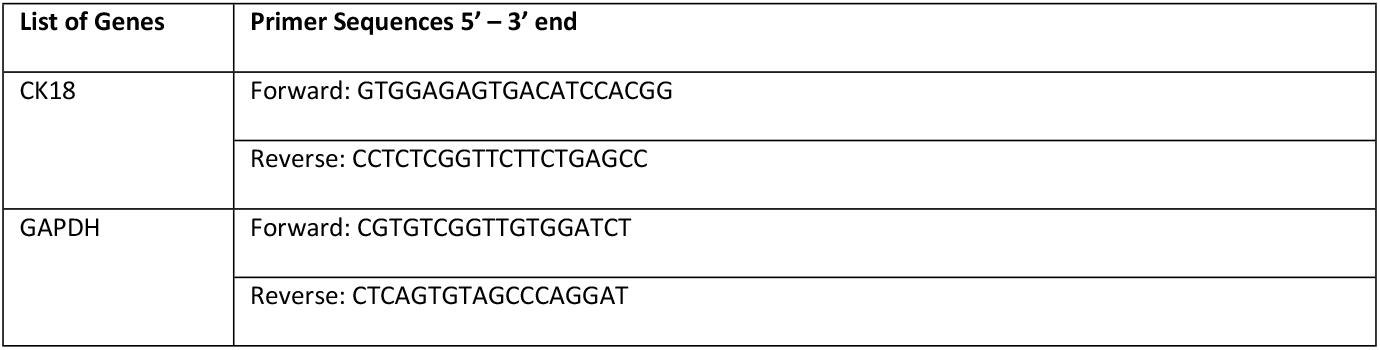
List of genes and primer sequences

